# Detection of horizontal sequence transfer in microorganisms in the genomic era

**DOI:** 10.1101/2022.12.21.521446

**Authors:** Mamadou Lamine Tall, Maxime Descartes Mbogning, Edmond Kuete Yimagou, Didier Raoult, Anthony Levasseur

## Abstract

Sequence transfer and genome remodeling are very frequent events in microorganisms, especially prokaryotes. This is due to the mosaic structure of the genomes, which calls into question the correct classification of genomes in terms of a single gene or a group of genes. We started here, as a first step, the inventory of bioinformatics tools applied to the detection of horizontal sequence transfers in microorganisms in the genomic era by applying bibliometric survey. Our bibliographic analysis allowed us to identify 17 main tools for the detection of chimeras. The first Bellerophon developed in 2004, followed by CCode and Pintail in 2005; Mallard in 2006; Mothur in 2009; Blackbox Chimera in 2010; Perseus, ChimeraSlayer, UCHIME 1 and UCHIME2, and ChimeraScan in 2011; Decipher in 2012; EBARDenovo and FunFrame in 2013; CATCh in 2015; Uchime 2 in 2016 and ChimeraMiner in April 2019. We then described each of these tools, highlighting their operating principles as well as the advantages and limitations of each (specificities, sensitivity, rapidity and frequent updates). The number of articles citing these tools has increased over the years especially for Mothur, Uchime, VSEARCH and ChimeraSlayer, thus demonstrating the interest of researchers in these tools and the need to decipher chimeras in genomic era.

## INTRODUCTION

Exchanges of genetic material between different species were very common during evolution (Eisen 2000; Smith et al. 1992). Therefore, the detection of horizontal transfer was considered crucial for the understanding of evolutionary processes and for the qualitative and quantitative assessment of the exchange rate between microorganisms (Dutta & Pan 2002; Syvanen 1994). The advent of new sequencing technologies has opened new avenues for evolutionary biology research. The sequencing of new microbial clades and the depth of sequencing, provided by the new sequencers, make it possible to study in depth the mechanisms of evolution of these genomes. In particular, the mosaicity of genomes has been highlighted and the origin of genomic content has been decrypted via the generation of “evolutionary bushes” or rhizomes(Levasseur et al. 2017). This approach consists in integrating all the coding sequences and determining their origin (HGT detection, horizontal gene transfer). Moreover, mosaic structures within the sequences could also be highlighted (horizontal sequence transfer). Beyond gene transfer, the transfer of sequences thus calls into question the current criteria of taxonomy. Indeed, the use of individual molecular markers for taxonomy does not take into account the mosaicity of exploited sequences (Haas, Gevers, Earl, Feldgarden, Ward, Giannoukos, Ciulla, Tabbaa, Highlander, Sodergren, Methé, DeSantis, Petrosino, et al. 2011). In addition, the use of several concatenated markers further biases the phylogenetic signal. To do so, we carried out a state-of-the-art study of applied bioinformatics, the detection of horizontal sequence transfers in microorganisms in the genomic era. Thus, we define in this section the needs for sequence transfer analysis tools, from sequence quality control through pre-processing to the analysis of genomic sequences by applying bibliometric survey. Then, we describe the limitations and advantages for each tool. We list the tools and web resources available to perform sequence transfer analysis. The global trend of research on chimera detection tools is also introduced. Finally, their consequences in metagenomics, evolutionary biology and taxonomy are discussed.

## MATERIALS AND METHODS

### Tool selection and data collection

Information from databases of scientific publications (such as NCBI PubMed, Web of Sciences (WOS)) has been filtered to allow us to find recent tools, software or software packages that aim to verify chimeras A keyword search for chimera check (“Chimeric Read” OR “Chimeric data” OR “chimera detection” OR “detect chimeric sequences” OR “chimera finder” OR “chimera check”) was used. The search yielded from 1354 articles in PubMed and Medline, 52 articles from WOS on 05/07/2019. Subsequently, these articles were screened manually by reading title and abstract. Only these 17 papers, the first papers describing new tools for chimera control, have been selected and studied in detail. The total records of those publications, including years of publications, titles, names of authors, affiliations, nationalities and names of publishing journals were collected from each publication. Information on the tools and their operating principles was collected from the original paper. The modeling of the operating principle of the tools was carried out using the draw.io software (https://www.draw.io/). Excel was used to create all the graphs in this article.

### Bibliometric analysis and Visualization

Using the WOS database, we collected the articles citing the original papers for different chimera tools. Total citations counts were collected from the Web of Science database on 06/07/2019. The lists of papers citing the different tools were exported to Excel 2016. They were subsequently merged, and the duplicates were removed. We used VOS viewer (Leiden University, Leiden, Leiden, The Netherlands) for constructing and visualizing bibliometric networks(Synnestvedt et al. 2005) (http://www.vosviewer.com).

## BIOINFORMATICS AND STATISTICAL TOOLS FOR CHIMERA DETECTION

Our bibliographic analysis allowed us to identify 17 tools for the detection of chimeras. The first Bellerophon developed in 2004, followed by CCode and Pintail in 2005; Mallard in 2006; Mothur in 2009; Blackbox Chimera in 2010; Perseus, ChimeraSlayer, UCHIME 1 and UCHIME2, and ChimeraScan in 2011; Decipher in 2012; EBARDenovo and FunFrame in 2013; CATCh in 2015; Uchime 2 in 2016 and ChimeraMiner in April 2019. Figure 1 illustrates this dynamic in the development of these tools. We will describe each of these tools, highlighting their operating principles and the advantages and limitations of each.

**Figure1:**
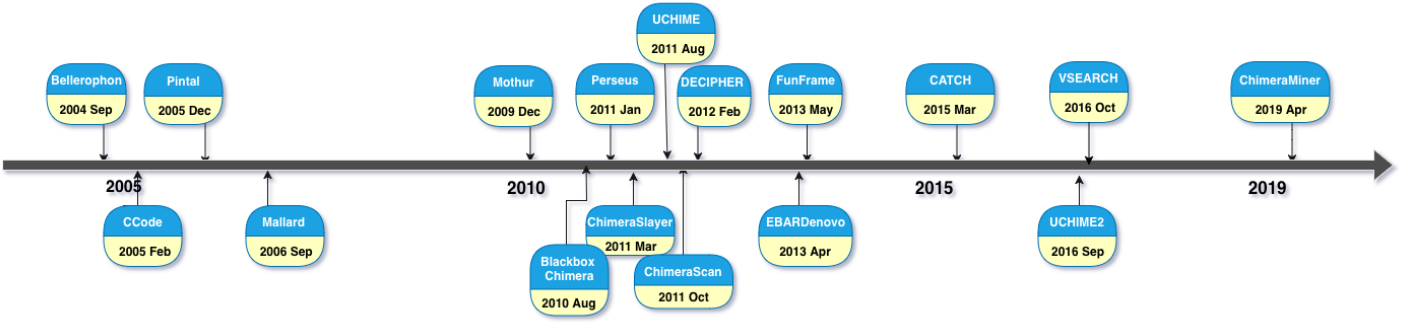
Modeling Chimera detection tools

### 1. Bellerophon

Bellerophon (Huber et al. 2004a) is a tool that also allows the detection of chimeric sequences specific to the 16S rRNA genes developed by Greengenes (DeSantis, Hugenholtz, Larsen, et al. 2006a) through several sequence datasets. It is based on a partial tree approach (Wang & Wang 1997), i.e. the phylogenetic tree that is derived from the independent regions is the result of a multiple alignment whose branching patterns are compared to indicate chimeric sequences.

### 2. CCode

To perform an analysis of suspected chimeric sequences, it is necessary to have all the closest sequences in a database. A comparison will be made between the request sequences and the reference sequences in order to confirm to confirm or refute the existence of a chimeric origin. Note that a chimeric sequence is composed of at least two partial gene sequences and each fragment is examined according to size and distance. The precision on the chimeric or non-chimeric origin of a query sequence is based on a 95% confidence threshold value and an analysis of variance test (Sokal Rolph). CCode (Gonzalez et al. 2005) (Detection and evaluation of chimera and crossing) is written in programming language C available for free at (http://www.rtphc.csic.es/download.html and on http://www.irnase.csic.es/users/jmgrau/index.html). Note that the reference sequences used in this program were obtained using the blastn algorithm (Altschul et al. 1990). Using too long or too short, a sequence can cause a resignation of of the program sensitivity.

### 3. Pintial

Pintial(Ashelford et al. 2005a) is a utility for the detection and analysis of 16S rRNA chimeras and other abnormalities. The operating principle is very simple as it uses a graphical interface and is capable of running on various computer platforms. It compares the length evolution distance of the 16S rRNA gene that exists between the query sequence and the subject sequence by advancing a fixed number of bases at a time along the length of the gene. Using this method, 1399 sequences of 19 phyla, as defined by the RDP project, were sorted and found that 64.3 were chimeras, 14.3% were non-identifying sequencing errors and 21.4% were strongly degenerate. Although Pintial was originally a 16S anomaly detection tool, most of the anomalies it detects are chimeras.

### 4. Mallard

Mallard (Ashelford et al. 2006a) is a computer-based tool for the detection of chimeras and other artifacts written in Java 1.4 programming language based on 16S rRNA genes. Mallard develops the Pintial algorithm [15] which runs by making a pairwise comparison between the reference sequence and the query sequence based on an alignment to evaluate changes in distance that are scalable and uncorrected. The outlier values obtained are identified as those that are below the threshold set by the user for the sequence comparison. Mallard was used in the analysis of the rRNA gene database with the same principle, i.e a multiple sequence alignment with the reference sequence in order to detect putative abnormalities that are verified with BLASTn. The comparison of Mallard with Bellerophon (Huber et al. 2004b) indicates show that on average 73.1% of chimeras know were by Mallard counter to Bellerophon which on average 59.8% of the chimeras were detected. Mallard more successfully detects putative abnormalities in bacterial and archaic taxons.

### 5. Mothur

Mothur (Schloss et al. 2009a) both to be a software package that brings together all the software intended for the microbial community, including the chimera detection tools (ChimeraSlayer and Uchime) in the form of a single tool capable of analyzing all these data. Mothur is written in C ++ with the help of object-oriented programming strategy. The first version was developed in 2009 and implements a lot of analysis algorithms for the 16S rRNA gene. The design patterns used in this program are enhanced for flexibility and software maintenance. This during mothur has a limit on the speed of the RAM available on the user’s computer.

### 6. Blackbox Chimera check (B2C2)

Blackbox Chimera check (Gontcharova et al. 2010). Chimera check is a chimera filtering tool that is based on a graphical interface executable in the Windows NET platform to process a batch of sequence of wide range. During the same process, several files initially selected can be analyzed. The regions at the beginning and at the end of the contigues from the input sequences are aligned through a BLAST database.

### 7. Perseus

The Perseus algorithm exploits the abundance of the sequences that are associated with the pyrosequencing data by considering each sequence at a time by performing precise pairwise alignments on all sequences of equal or greater abundance, on all possible parents (Quince et al. 2011a). A query is marked as chimera if the distance calculated during the alignment is less than 0.15 and less than or equal to the distance of the closest sequence between the best possible parents. This hypothesis makes it possible to validate the existence of a chimera. Taking into account that this chimeric model may have evolved, another factor called “chimera index” will be calculated based on the query, the closest relative and the parent farthest away using ancestral parsimony. Once the index is determined, there remains the supervised learning through a set of data sets (V2 and V5) or for each sequence, a chimera matching criterion will be calculated, then a classification will be performed to determine by comparison with the reference sequences to form a logistic regression.

### 8. Algorithm ChimeraSlayer CS

The algorithm (Figure 2) runs by first passing through a NAST (DeSantis, Hugenholtz, Keller, et al. 2006) format alignment using the NAST-iEr multiple alignment utility. Using this output file, the end regions of the request sequence each corresponding to 30% of the request length will be independently searched in the 16S reference database using Megablast. An identification of the chimeric parent will be made, so that it is an in-silico chimera among several parent reference 16S having a high alignment score. The best results for each search will be extracted in NAST format through a dynamic alignment program CHECK-CHIMERA (Komatsoulis & Waterman 1997). After identification, another NAST realignment will be done, but this time based on the profile of the selected parents. Finally, the query sequences, which present a great similarity with an in silico chimera formed between any two parental sequences, are marked in an evolutionary framework, i.e. a bootstrap greater than 90% and a minimum divergence greater than 1.007 (Haas, Gevers, Earl, Feldgarden, Ward, Giannoukos, Ciulla, Tabbaa, Highlander, Sodergren, Methé, DeSantis, Consortium, et al. 2011a; Komatsoulis & Waterman 1997).

**Figure 2:**
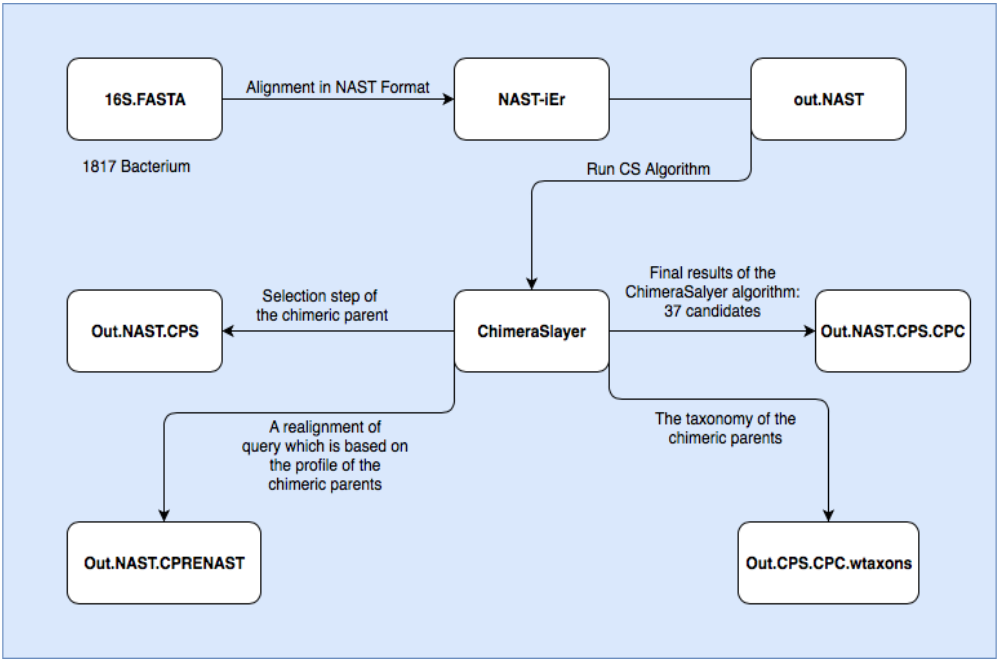
Modeling ChimeraSlayer Algorithm.

### 9. Uchime

Uchime (Figure 3) uses the query sequence and divides it into four pieces or segments, each portion is searched through a reference database, and the best matches of each search are noted. Among these correspondences, the two best candidates are identified, and a multiple alignment will be done (local, local-X and global-X) based on the profile of the two selected candidates. If the percentage sequence identity is ≥ 0.8%, an alignment score will be calculated and if the threshold is exceeded, the query is marked as chimera(Edgar et al. 2011a).

**Figure 3:**
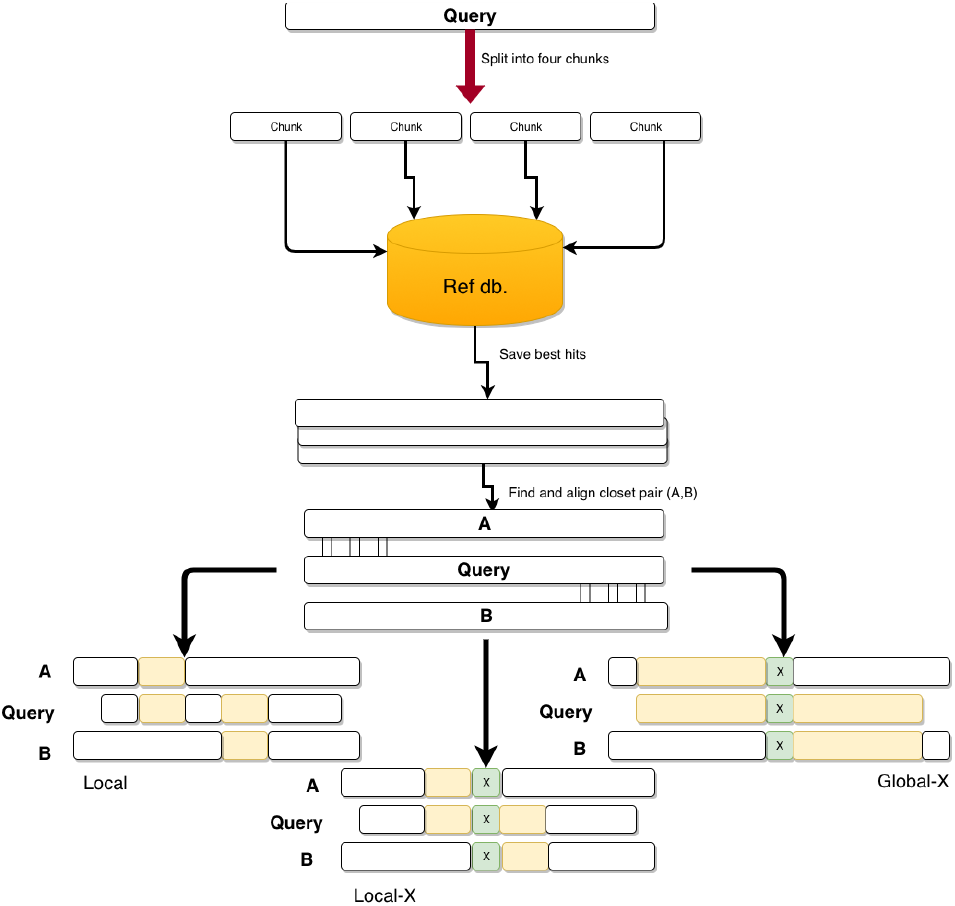
Modeling Uchime Algorithm: The query sequence is divided into 4 pieces; each piece is searched in the reference database and the best ones are selected after calculating the percentage of identity. Three types of alignment are made (Local, local-X and Global-X) between the query (Q) and both parents (A, B)(Edgar et al. 2011a).

### 10. ChimeraScan

ChimeraScan is an open source chimera detection software package that includes features such as: processing long matched reads greater than 75bp, handling ambiguous mapping reads, detecting readings covering a merge junction, integrating with very popular aligner of nodes (Langmead et al. 2009). The comparison of ChimeraScan with other tools, such as deFuse (McPherson et al. 2011), shortFuse (Kinsella et al. 2011) and MapSplice (Wang et al. 2010) shows that 90% of the positives among 78 chimera predictions were detected by ChimeraScan.

### 11. Decipher

DECIPHER(Wright et al. 2012) is a new method for detecting chimeric 16S rRNA sequences using a chimeric sequence search approach. It is based on the detection of short fragments that are rare in a phylogenetic group where a query sequence is classified but found in another phylogenetic group. Calibration of the algorithm was carried out for complete sequences (fs_DECIPHER) and short sequences (ss_DECIPHER) and was then compared to other algorithms namely WigeoN (Pintail) (Ashelford et al. 2005a), ChimeraSlayer (Haas, Gevers, Earl, Feldgarden, Ward, Giannoukos, Ciulla, Tabbaa, Highlander, Sodergren, Methé, DeSantis, Consortium, et al. 2011b) and Uchime (Edgar et al. 2011a). In fact ss_DECIPHER and Uchime provided a better detection of chimera on short sequences of around 100 to 600 nucleotides in length, ie 79% and 81%, but if we move on to longer sequences, Uchime’s performance begins to deteriorate while those of ss_DECIPHER increases with a detection rate around 89%. Based on either tool, positive rates were estimated to be approximately 1.3% and 1.6%. Indeed, the rate of false positive of fs_DECIPHER was very low compared to other programs, i.e. a rate of less than 0.20%. DECIPHER was only surpassed by ChimeraSlayer and Uchime when the chimeras are formed from close relatives, a difference of less than 10%. With fs_DECIPHER, the detection of chimeras in the RDP (Cole et al. 2007), SILVA(Pruesse et al. 2007) and Greengenes(DeSantis, Hugenholtz, Larsen, et al. 2006b) were outperformed, equivalent to 1% and 2% more chimeras.

### 12. EBARDenovo

EBARDenovo (Chu et al. 2013) runs in three main stages: extension, bridge and chimera detection. Before performing these steps, the algorithm points to the first base pairs for each read (default 15pb) as the index key, then a sort will be applied based on read abundance. As part of the detection of sequence repeat detection (chimeras), predictable readings are used to identify repetitive sequences. EBARDEnovo is implemented in C programming language and can be run on Windows, Linux and Windows operating systems. 64-bit Mac OS. It is imperative to install the Mono program for execution (http://www.mono-project.com/).

### 13. Funframe

FunFrame is a pyrosequencing error analysis pipeline using HMM-FRAME (Zhang & Sun 2011) ; UCHIME (Edgar et al. 2011a) for chimera detection and ESPRIT-Tree (Cai & Sun 2011) for clustering using the OTU array. FunFrame is coded in R and Python programming language and the execution is performed by a bash script and can be done individually.

### 14. CATCH

CATCH (Mysara et al. 2015), which groups the classifiers (CATCH reference and CATCH de novo) is a machine learning algorithm for the detection of chimeric and non-chimeric sequences from a set of input data. Using CATCH, the input data are scores from the various chimera detection tools such as mothur (Schloss et al. 2009b) that implement the algorithms (ChimeraSlayer UCHIME and Pintial) that use the Gold Reference Database (on the Broad Microbiome Utilities website, version micorbiomeutil-r20110519) and DECIPHER that uses the RDP. This database during each tool is run separately and the output scores are processed by a classifier to finally predict whether the reading is chimeric or not. This automatic learning is done in three steps: (i) firstly, the identification of the output values of the different tools for detecting chimeras, (ii) in a second step, the classifier is trained in a supervised learning it learns to establish a correct prediction according to the given function from the tools and, at the end, (iii) the learning is validated the classifier is currently being able to predict chimeric sequences with other data different from those used as examples. For the reference-based CATCH classifier, the features are (i) the score and final decision that are calculated for UCHIME and ChimeraSlayer (ii) and the standard deviation and final decision that are calculated for Pintial and (iii) the final decision that remains for DECIPHER. While for the de novo CATCH classifier, the score characteristics and final decision are chosen as input parameters for UCHIME, ChimeraSlayer and Perseus.

### 15. Uchime2

UCHIME2 (Edgar 2016) is an optimized update of the UCHIME (Edgar et al. 2011b) chimera detection algorithm that operates by building a model and performing multiple alignment of several models and the most important results of the most similar sequence. If the fractional divergence is large, then one has a model and the query is more likely to be marked as a chimera and if the divergence is small, the model is more likely to be a false.

### 16. VSEARCH

The VSEARCH algorithm (Rognes et al. 2016) uses a fast heuristic based on databases shared by the query and target sequences to identify similarities between them. This strategy is also used by USEARCH (https://www.drive5.com/usearch/). With VSEARCH, dynamic programming will perform optimal global alignment of the query on potential target sequences and pairwise alignments are calculated through vectorization and multiple threads. It is a multithreaded 64-bit tool that is free and open source and intended for metagenomic data analysis.

### 17. ChimeraMiner

ChimeraMiner (Lu et al. 2019a) is an improved chimera detection tool for the analysis of MDA (Multiple Displacement Amplification) sequencing data. To compare the efficiency of the algorithm with the previous tools, two sets of MDA data sets were used (MDA1 and MDA2). In terms of treatment time, it is estimated at 43.4% and that 83.60% of the chimeras detected by the previous tools were found by ChimeraMiner, plus 6,736,168 chimeras not detected by the previous methods.

## GLOBAL TREND OF CHIMERAS TOOLS RESEARCH: A BIBLIOMETRIC ANALYSIS

Bibliometrics applies mathematical and statistical techniques to the analysis of books, articles and other documents (Otlet 1934; Pritchard 1969), thus highlighting the current level of knowledge in a scientific field through the compilation of data obtained from bibliographic databases and their quantitative and qualitative analysis (Ekinci et al. 2015). Bibliometrics can be used to inform policy decisions (Russell & Rousseau 2010), allocate research funding (Cronin & Sugimoto 2014), help libraries prioritize acquisitions(Engler 2014) and evaluate scholarly activities (Rousseau et al. 2018). The objectives of this section were to provide a brief bibliometric analysis of the articles that cited the original articles from each of these tools.

### Bibliographic analysis of chimera’s detection tools original papers

The authors who contributed to the development of these tools came from 13 different countries. The USA contributed to the development of these 10 tools (Uchime et Uchime2, FunFrame, DECIPHER, ChimeraScan, Chimera Slayer, Blackbox Chimera check (B2C2), Perseus, Bellerophon et Mothur) ; 06 from United Kingdom (Perseus, Mallard, Uchime, Mothur, VSEARCH and Pintail) ; 02 from Canada (Mothur and EBARDenovo) ; 02 from Norway (VSEARCH and Perseus). The authors from Taiwan, Spain, Belgium, China, Australia, Germany, France, Slovenia and Austria contributed respectively for EBARDenovo, CCode, CATCh, ChimeraMiner, Bellerophon, VSEARCH, VSEARCH, Mothur and Mothur (Table 1). Bioinformatics is the journal in which most of these papers (06 publications) were published, followed by Applied and Environmental Microbiology (05 publications) and one in each of the bioRxiv, BMC Bioinformatics, Genome Research, International Journal of Molecular Sciences, PEERJ and Open Microbiology Journal (Table 1). Among the features of chimera detection tools, the most commonly used programming language is C++, the reference interface is often command line based and stability is checked for most tools (Table 2).

**Table 1:**
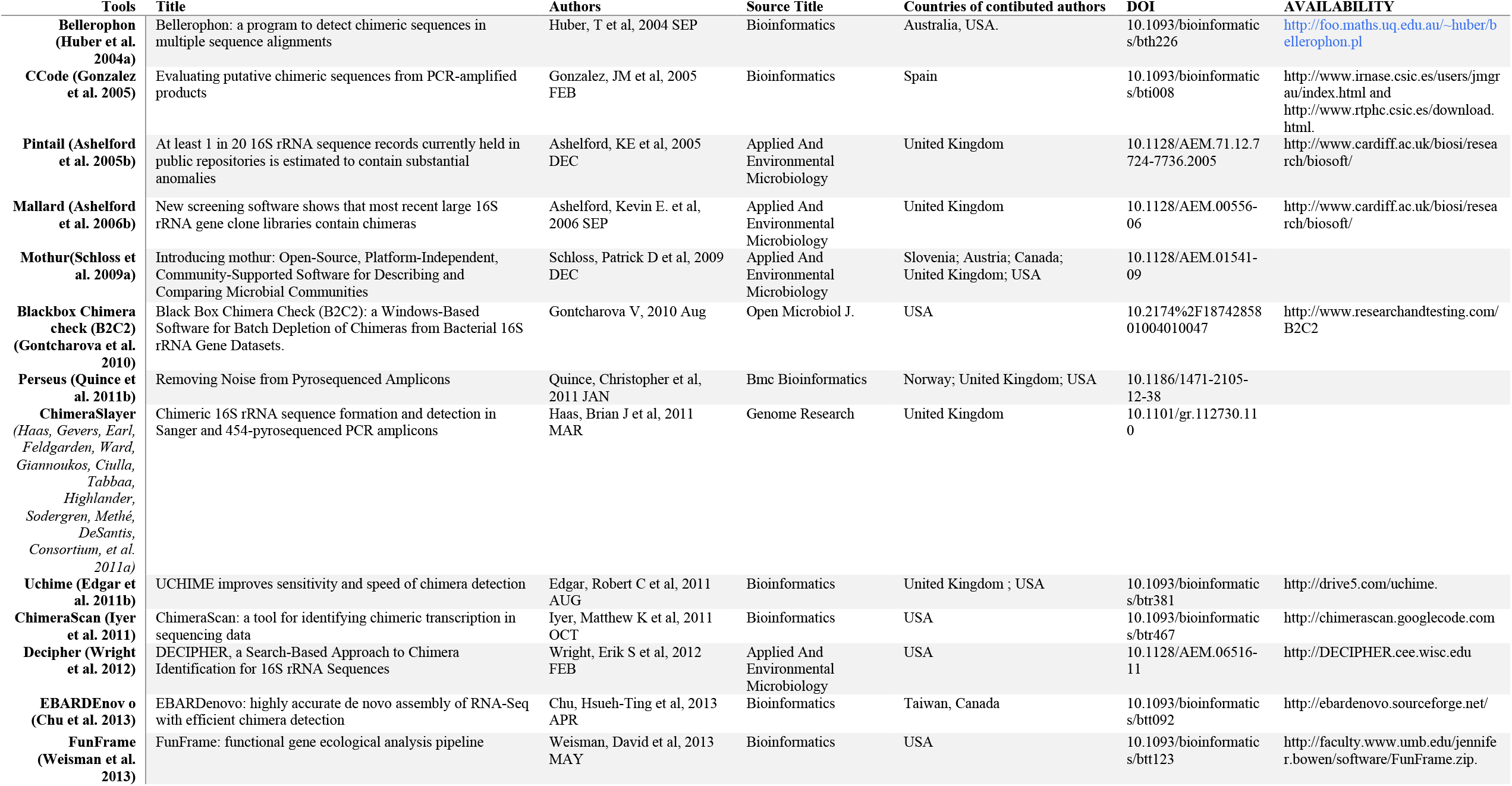
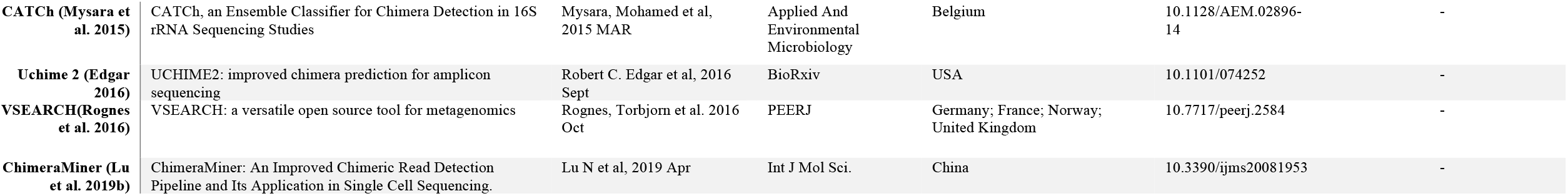
Bibliographic Informations of chimeras detection tools.

**Table 2:** Characteristics of chimera detection tools.

| Tools | Software type | Interface | Restrictions to use | Biological technology | Implementation | Operating system | Computer skills | Stability | Maintained | Version |
| --- | --- | --- | --- | --- | --- | --- | --- | --- | --- | --- |
| **Bellerophon(Huber et al. 2004a)** | Pipeline/ Workflow | Command line interface | None |  | Python | Unix/Linux | Advanced | Stable | Yes | - |
| **CCode (Gonzalez et al. 2005)** | - | - | - |  | C |  | - | - | - | - |
| **Pintail (Ashelford et al. 2005b)** | Achievable | Graphical interface |  |  | Java | Linux, Windows et OS X | Advanced | Stable | Yes | 10.2 |
| **Mallard (Ashelford et al. 2006b)** |  |  |  |  | Java |  |  |  |  | - |
| **Mothur (Schloss et al. 2009a)** | Package/ Module | Command line interface | None |  | C++ | Unix/Linux | Advanced | Stable | Yes | 1.38.0 |
| **Blackbox Chimera check (B2C2) (Gontcharova et al. 2010)** | Achievable | Graphical interface |  | Roche | C++ | Linux/Windows | Advanced | Stable | Yes | 0.62.x |
| **Perseus (Quince et al. 2011b)** |  |  |  |  |  |  |  |  |  |  |
| **ChimeraSlayer (Haas, Gevers, Earl, Feldgarden, Ward, Giannoukos, Ciulla, Tabbaa, Highlander, Sodergren, Methé, DeSantis, Consortium, et al. 2011a)** | Package/ Module | Command line interface | None | Roche | Perl | Unix/Linux | Advanced | Stable | Yes | - |
| **Uchime (Edgar et al. 2011b)** | Package/ modules | Command line interface | None |  | C++ | Unix/Linux | Advanced | Stable |  | 4.2.40 |
| **ChimeraScan (Iyer et al. 2011)** | Package/ Module | Command line interface | Academic or non-commercial use |  | Python | Unix/Linux | Advanced | Stable | Yes | 0.4.5 |
| **Decipher (Wright et al. 2012)** | Package/Module | R software | None |  | R programming language | Unix/Linux, Mac OS, Windows | Advanced |  | Yes | - |
| **EBARDe novo (Chu et al. 2013)** | Application/Script, Package/Module | Command line interface | Academic or non-commercial use |  | C | Unix/Linux, Mac OS, Windows | Advanced | Stable | Yes | 2.0.1 |
| **FunFrame (Weisman et al. 2013)** | Pipeline/ Workflow | Command line interface | None |  | R | Unix/Linux, Mac OS, Windows | Advanced | Stable | Yes | 0.9.3 |
| **CATCH (Mysara et al. 2015)** | Package/Module | Command line interface | None | Illumina, Roche | - | Unix/Linux | Advanced | Stable | Yes | - |
| **UCHIME2 (Edgar 2016)** | Package/Module | Command line interface | None |  | C++ | Unix/Linux | Advanced | Stable | Yes |  |
| **VSEARCH** | Package/Module | Command line interface | None |  | C, C++ | Unix/Linux, Mac OS, Windows | Advanced | Stable | Yes | 2.3.4 |
| **ChimeraMiner (Lu et al. 2019b)** | Package/Module | Command line interface | None |  | Perl | Unix/Linux, Mac OS | Advanced | Stable | Yes | - |

### Citation analysis of papers citing chimera’s detection tools

Of the 17 tools, only the 15 indexed in the WOS database were used for further analysis. These are: Mothur, UCHIME, Chimera Slayer, Bellerophon, Perseus, VSEARCH, Pintail, Mallard, DECIPHER, ChimeraScan, CCode, EBARDenovo, CATCh, FunFrame et ChimeraMiner. They were cited respectively 8424; 5121; 1335; 1246; 857; 590; 575; 552; 345; 141; 60; 15; 12; 8 and 0 times. And an average per year respectively of 765,82; 569; 148,33; 77,88; 95,22; 147,5; 38,33; 39,43; 43,13; 15,67; 4; 2,14; 2,4; 1,14 et 0 (Table 3). The total number of papers citing the chimera detection tools was 14876. The number of papers citing these tools has grown over the years for Mothur, Uchime, VSEARCH et Chimera Slayer which demonstrates the interest of researchers for these tools (Figure 4). Their specificity, precision, speed and frequent updates can be an explanation. There has been a loss of interest in tools such as Perseus, Pintail, Mallard, Decipher, ChimeraScan, CCode, EBARDenovo with the progressive decrease in the number of citations in recent years (Figure 4 and Figure 5).

**Table 3:** Number of citations of publications on chimera detection tools.

| Tools | Publication date | Total Citations | Average per Year |
| --- | --- | --- | --- |
| **Mothur** | Dec 2009 | 8424 | 765,82 |
| **UCHIME** | Aug 2011 | 5121 | 569 |
| **VSEARCH** | Oct 2016 | 590 | 147,5 |
| **Chimera Slayer** | Mar 2011 | 1335 | 148,33 |
| **Perseus** | Jan 2011 | 857 | 95,22 |
| **DECIPHER** | Feb 2012 | 345 | 43,13 |
| **Bellerophon** | Sep 2004 | 1246 | 77,88 |
| **Pintail** | Dec 2005 | 575 | 38,33 |
| **ChimeraScan** | Oct 2011 | 141 | 15,67 |
| **Mallard** | Sep 2006 | 552 | 39,43 |
| **CATCh** | Mar 2015 | 12 | 2,4 |
| **CCode** | Feb 2005 | 60 | 4 |
| **EBARDenovo** | Apr 2013 | 15 | 2,14 |
| **FunFrame** | May 2013 | 8 | 1,14 |
| **ChimeraMiner** | Apr 2019 | 0 | 0 |

**Figure 4:**
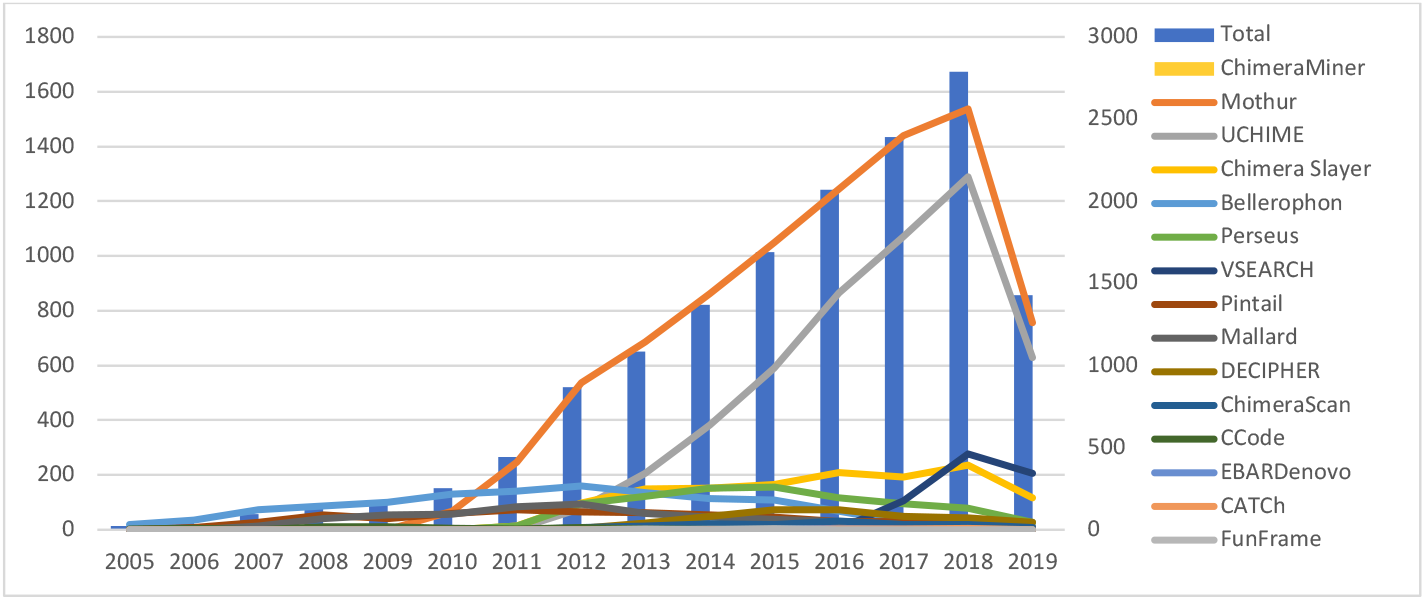
Evolution of the number of citations of chimera detection tools

**Figure 5:**
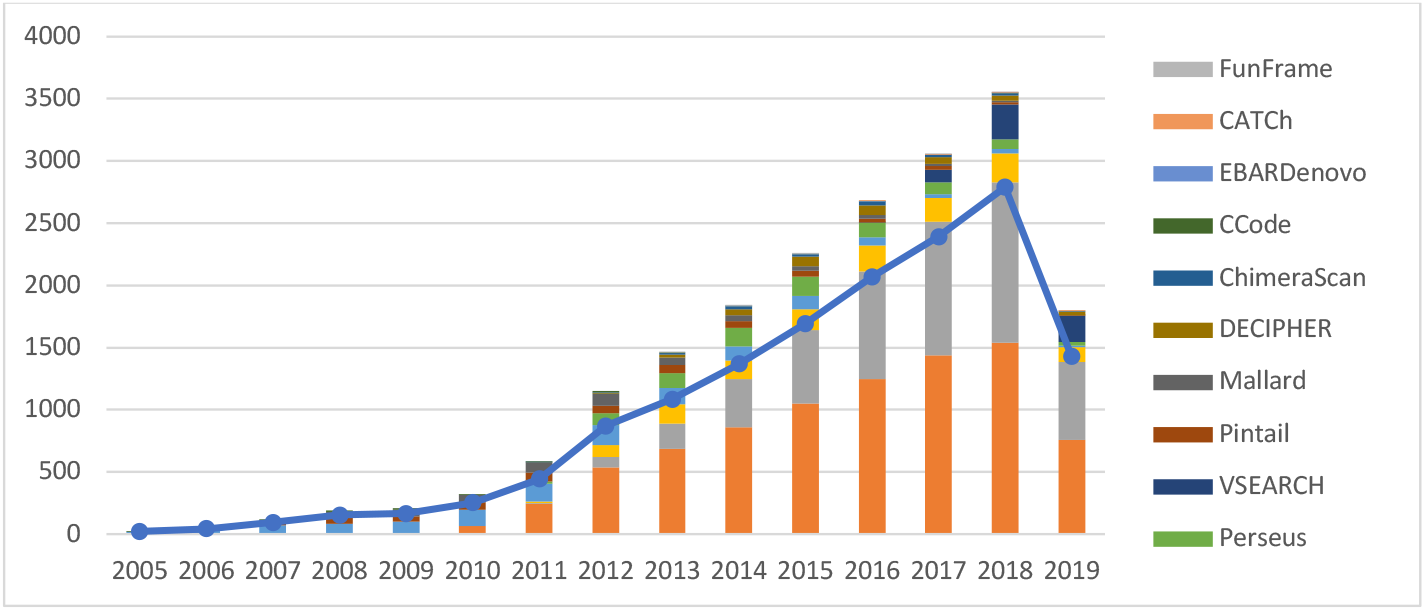
Evolutions curved of the top 20 most productive organizations

### Distribution of published journals on papers citing chimera’s detection tools

The citing document was published in 1525 journals, with 788 who published more than 2 citing documents. Plos One is the most used (1054 documents), followed by Frontiers in Microbiology (1013 documents) and Scientific Reports (608 documents) (Figure 6). After the year 2015, we can see in Figure 7 the decrease of the most used source (Plos One, Applied and Environmental Microbiology; FEMS Microbiology Ecology and Environmental microbiology) and the increase of some sources such as Frontiers in Microbiology and new sources like Scientific Reports, Science of the Total Environment, Microbiome, and Peerj. These most interesting sources are interested by microbial ecology in human and environment.

**Figure 6:**
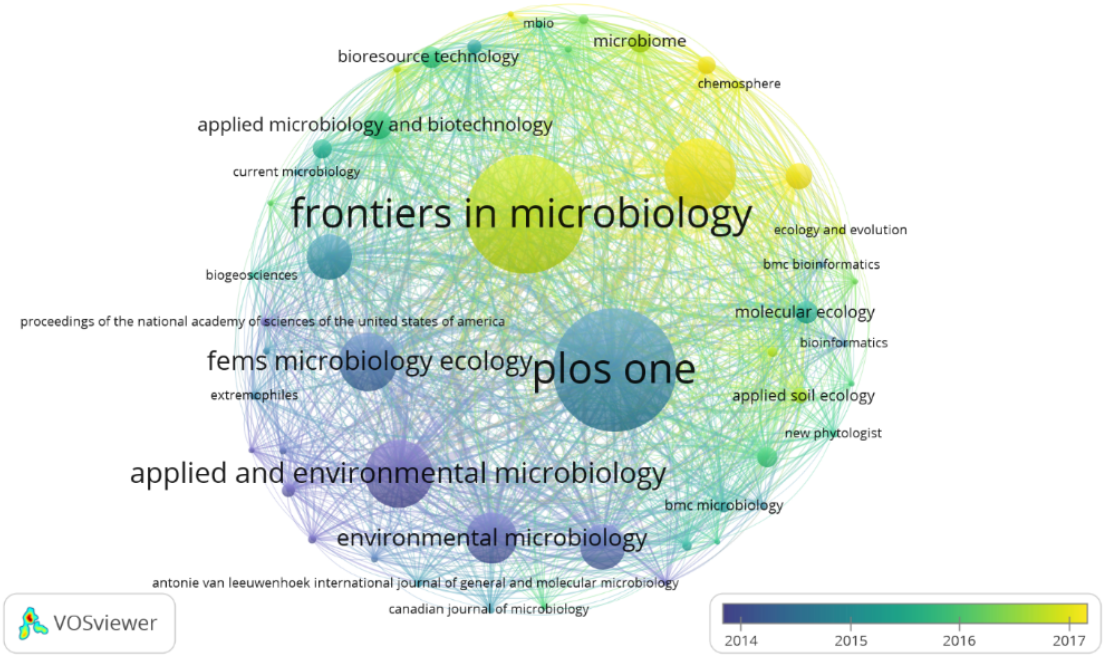
Combined mapping and clustering of top 50 most active journals according to the number of publications

**Figure 7:**
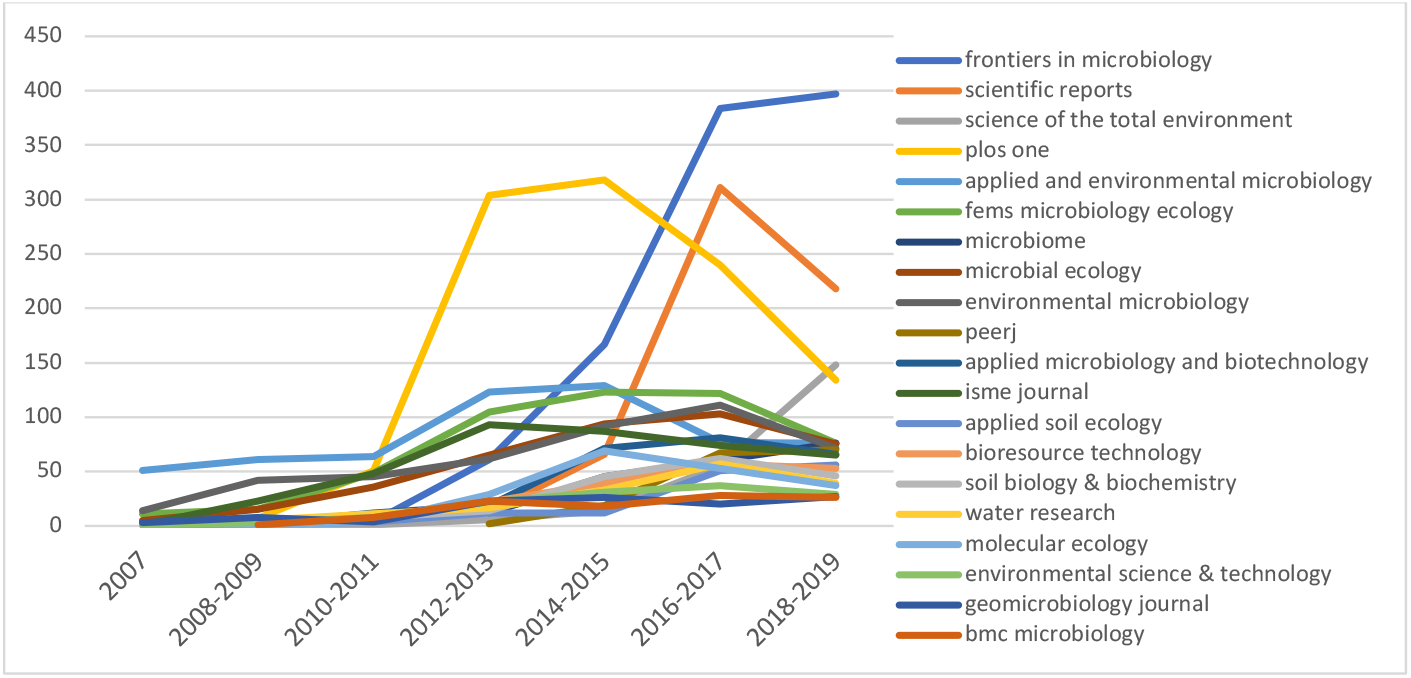
Evolutions curved of the top 20 most active journals according to the number of publications

### Countries contributing to global publications on papers citing chimera’s detection tools

More than 58,760 authors from various parts of the world have contributed to studies that have cited these chimera detection tools. In general, they were affiliated with institutions in 149 countries, the top ten of which was the USA first (5,146 items), followed by China (3,468 items), Germany (1,384 articles), Canada (967 articles), France (902 articles), Australia (811 items), England (804 items), Spain (741 items), Japan (598 items) and South Korea (555 items) (Figure 8). American documents were the most cited with 202,396 citations, followed by German documents with 44,309 citations and China with 42,219 citations. After 2017, China has become the country with the highest number of publications citing Chimera tools (Figure 9). Furthermore, they developed the last chimera tools to our knowledge, ChimeraMiner (Lu et al. 2019b), released early this year. France, a pioneer in this field, has also increased its position and is now one of the top 4 countries interested in studies related to the Chimera (Figure 9).

**Figure 8:**
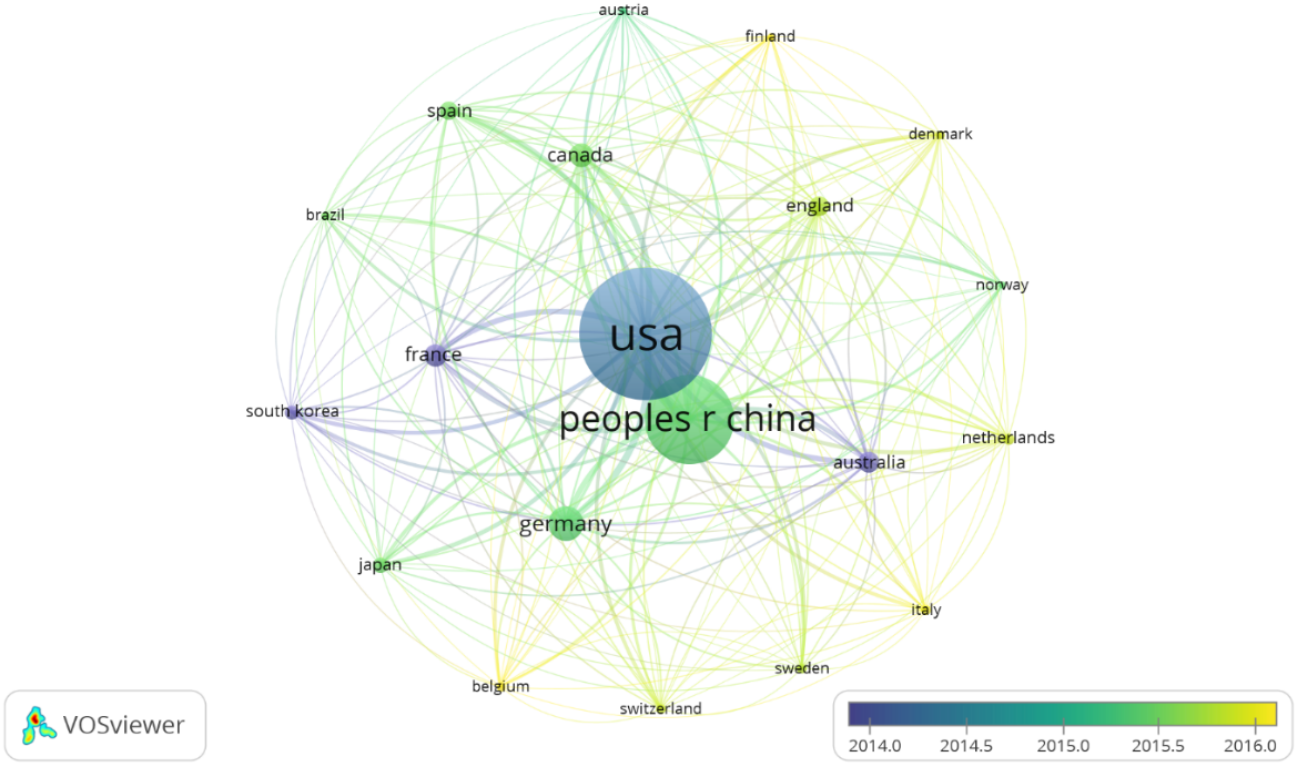
Networked visualization of collaborations between countries among the 20 most productive countries of publications citing chimera detection tools. Links represent the strength of collaboration

**Figure 9:**
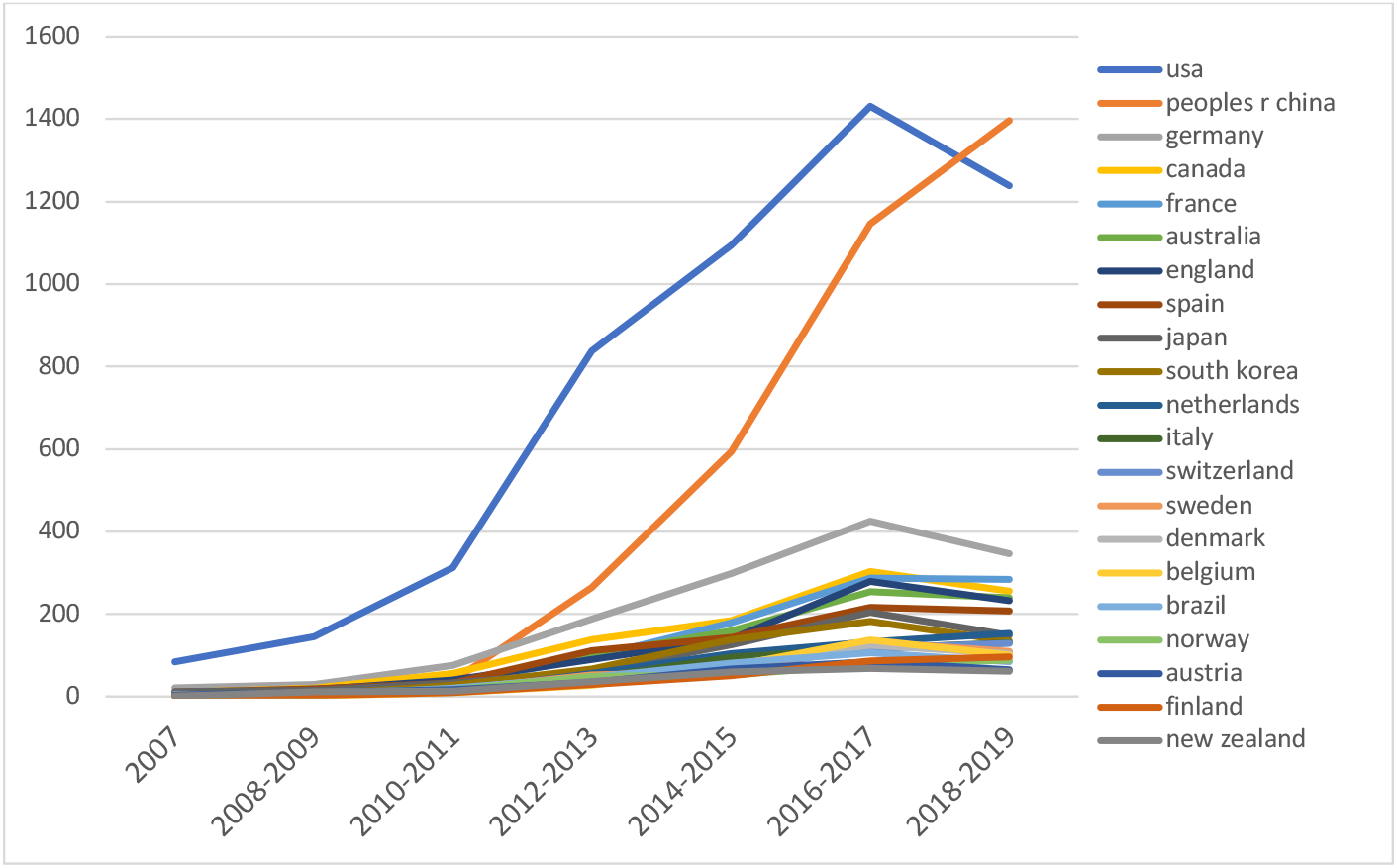
Curved evolution of the 20 most productive countries

### Distribution of institutes paying attention to chimera detection tools

The authors contributed in the document citing these chimera stools are affiliated with 9000 organizations, and 1466 organizations have at least 1466 documents. Among the top 20 organizations frequently citing these stools, Chinese organizations are in first and second place (respectively Chinese Acad Sci and Univ Chinese Acad Sci), followed by American organizations (Univ Michigan) (Figure 10; Figure 11). The French institutions, INRA and CNRS are respectively in 17th and 18th place, in terms of contribution cited (figure 10).

**Figure 10:**
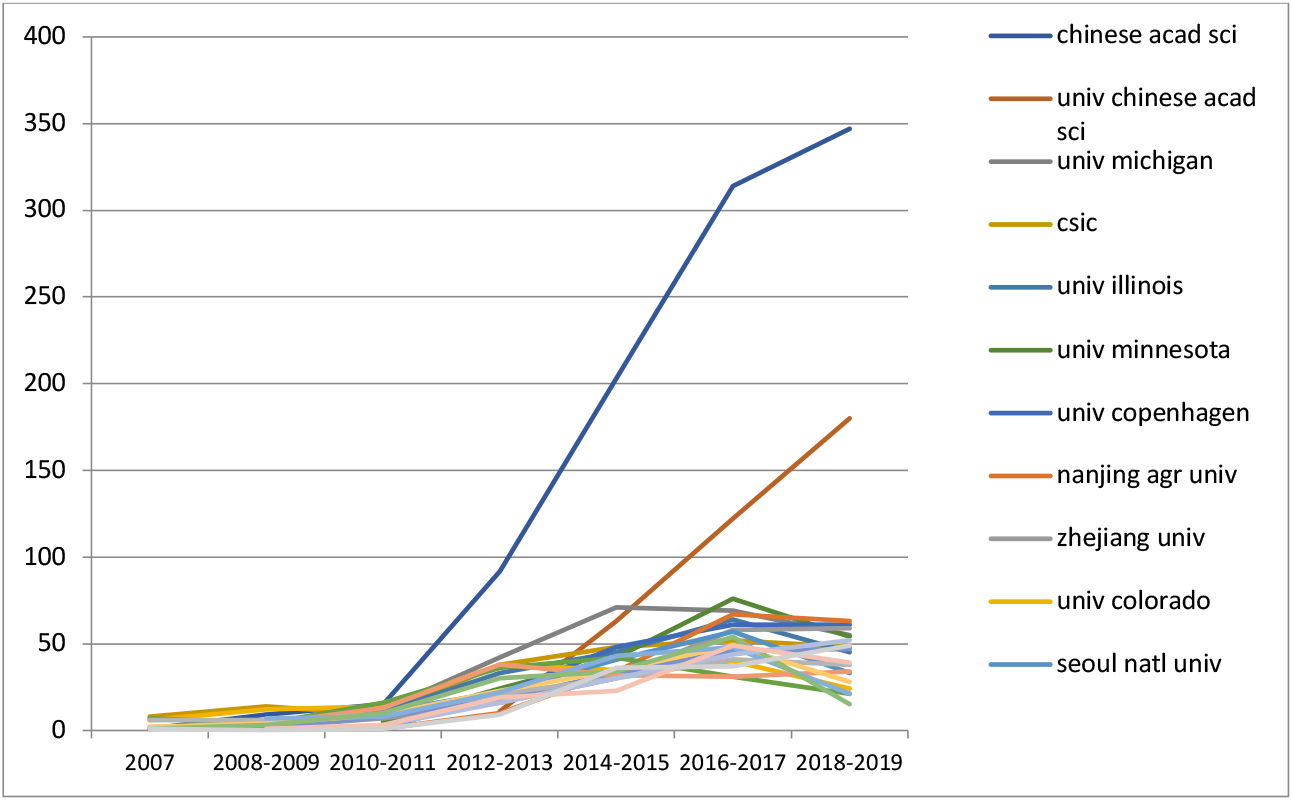
Curved evolution of the top 20 most productive organizations

**Figure 11:**
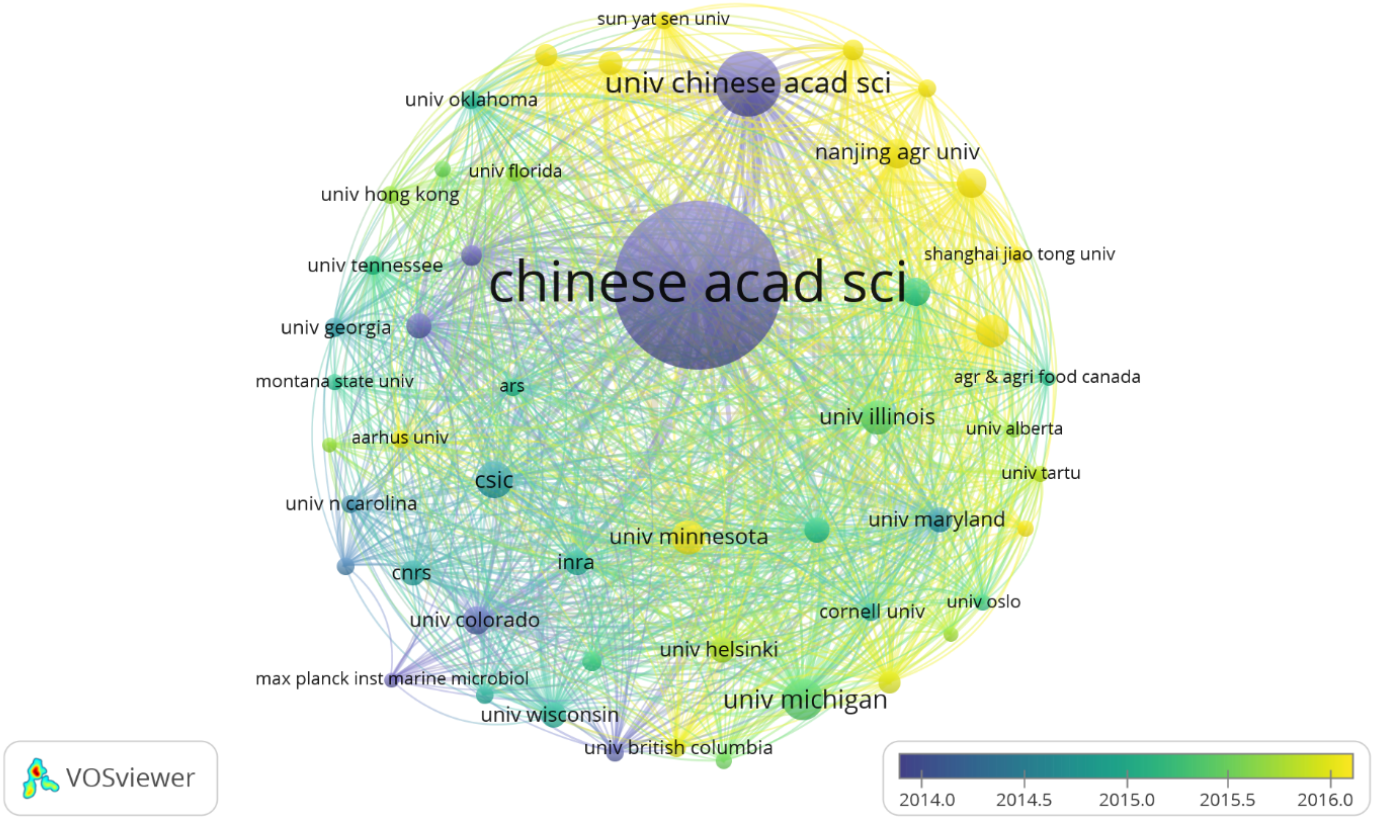
Networked visualization of inter-organizational collaborations among the 50 most productive organizations in publications citing chimera detection tools. Links represent the strength of collaboration

### Hotspots of studies citing chimera’s detection tools

We apply bibliometrics to the keywords of the analyzed articles. The keywords represent the author’s opinion (or about) the most important words of his articles. Second, keyword analysis can potentially detect trends in research topics, both currently and in the past (https://www.ncbi.nlm.nih.gov/pmc/articles/PMC6480778/). The authors of the citing document used 23670 keywords and 3646 were used more than 5 times. The 100 most frequent keywords used by the authors in the reference document were classified into 5 groups (Figure 12). The clusters are made up of the keywords related to microbiology techniques and microbial diversity. The group 1 color red, composed of 30 elements, is specifically constituted by the keywords related to wastewater treatment (activated sludge, wastewater, wastewater treatment and bioremediation) and biological processes induced by microorganisms (stable isotope probing, anammox, ammonia-oxidizing archaea, biodegradation, biofilm, denitrification and nitrification). Group 2, green in color, consisted of 29 items and the items specific to it are: bacteria (actinobacteria, bacteria and cyanobacteria), fungi (arbuscular mycorrhizal fungi, fungal community and fungi) and environment (Antarctica, bacterioplankton, rhizosphere, biogeography, arctic and soil). The Cluster 3, yellow in color and composed of 28 elements, is specifically made up of keywords related to human studies on probiotics, prebiotics, treatments (probiotics, lactobacilli and lactic bacteria, diet and antibiotics) and health status.(obesity, inflammation and dysbiosis). The group 4, of blue color and composed of 8 elements, is specifically constituted by the keyword relative to methanogenic archaea (anaerobic digestion, archaea, methane, methanogen, methanogenesis and methanogens). The last cluster 5, pink in color and composed of 6 elements, is specifically constituted by keywords relative to phylogeny and taxonomy (Figure 12, Figure 13). This analysis highlights the main research areas related to chimera stools. We can cite microbial diversity of human and environment; phylogeny and taxonomy; archaea and methanogen; human probiotics and dysbiosis; waste treatment and biological process induced by microorganisms.

**Figure 12:**
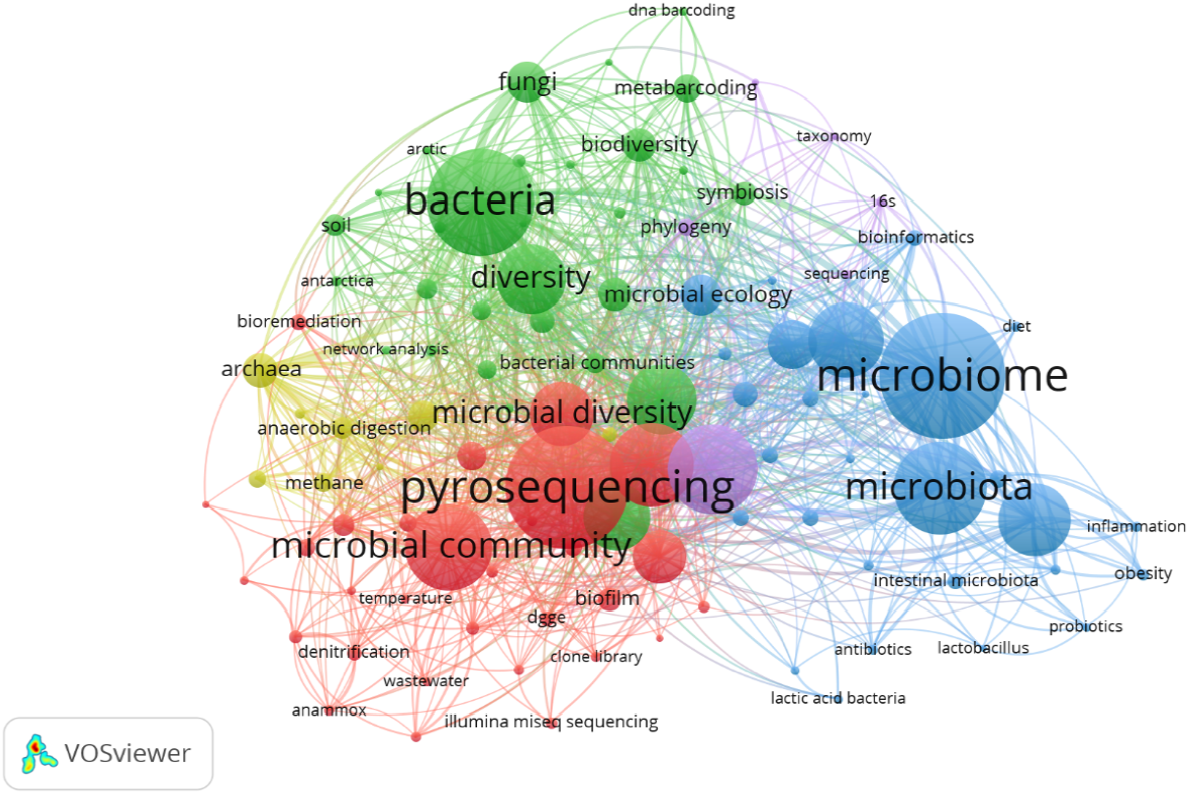
Network visualization of top 100 most used keywords of publications citing chimera’s detection tools. The keywords were divided into four clusters according to different colors generated using default paramaters: wastewater treatment (down in red), microorganisms and environment (up in green), archaea methanogen related study (right in blue), probiotics prebiotics and treatment related study (left in yellow) and phylogeny and taxonomy (up in pink). A large circle size represents the keyword that appears at a high frequency.

**Figure 13:**
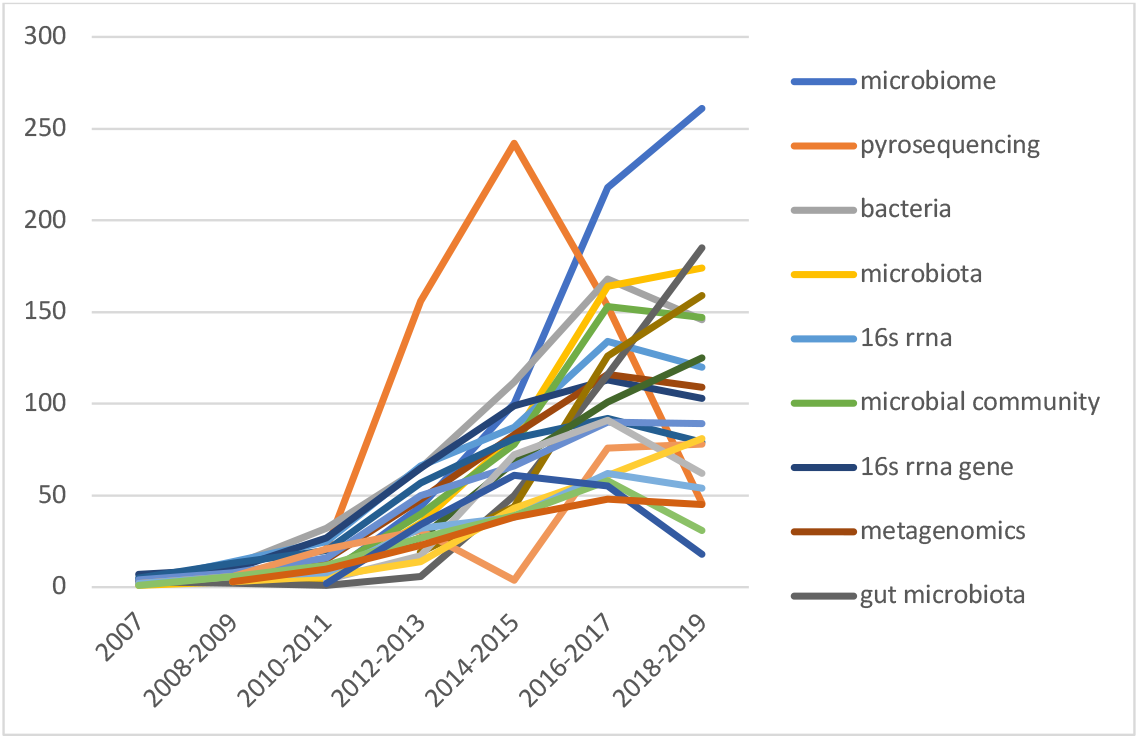
Evolutions curved of the top 20 most frequent used keywords

## THE CONSEQUENCES IN TERMS OF METAGENOMIC STUDIES AND EVOLUTIONARY BIOLOGY AND TAXONOMY

Advances in Next Generation Sequencing (NGS) have made a significant contribution to advanced microbial ecology studies, hence the implementation of metagenomics. This is defined as a process of analysis of the DNA without the need to perform a culture. With the implementation of databases and tools, bioinformatics and statistical analysis have made it possible to better exploit metagenomic data (Oulas et al. 2015). Together with marker genes, the use of metagenomics is an efficient approach to obtain a taxonomic profile of a given community using PCR amplification and sequencing of conserved marker genes such as 16S rRNA (Tringe & Hugenholtz 2008). The latter was considered a fundamental fold for the characterization of a given microbial community, but as a result, this approach is overshadowed by metagenomics, which provides the overall functioning of a community rather than an inventory of these inhabitants (Tringe & Hugenholtz 2008).

The essential function of the 16S gene, as well as its highly conserved sequence and structure, has made it the molecule of choice for microbial evolution and ecological studies [57,58]. The numerous highly conserved regions covering the length of the gene allow the amplification of sequences from a wide range of species. However, these same highly conserved regions contribute to cross hybridization and amplification priming errors that create chimeric sequences. The correct identification of chimeric 16S sequences is a difficult computational problem and they are more likely to be classified as novel organisms if not correctly identified as outliers.

## CONCLUSION AND DISCUSSION

The evaluation of the accuracy of sequence transfer detection for all the tools testifies to the great variety of algorithms. For example, BellerophonGG is able to identify chimeras mainly restricted to the most divergent sequence pairs, while Pintail is highly sensitive to the detection of chimeras in full-length sequences, not chimeras in shorter sequences.

However, the perfect detection of chimeras is still an unresolved problem. More complex chimeras and sequence anomalies may escape detection. This underlines the importance of obtaining and validating the sequences that represent the new bacterial diversity and of continuing to develop the reference database and additional analytical tools. The problem of identifying rare species that correlate with diseases or other important functions of the microbial ecosystem remains difficult. This method is the only one currently able to detect sensitive chimeras in short and long readings of 16S sequences. Indeed, perfect detection of chimeras is still an unresolved problem due to divergence and length, as detection accuracy begins to degrade as divergence from reference sequences increases.

## Notes

### Competing Interest Statement

The authors have declared no competing interest.

### Summary of Updates

I correct the authors affiliations

